# Climatic-environmental influences on hominin brain size over the last 5 million years

**DOI:** 10.1101/2024.09.09.611970

**Authors:** Samuel L. Nicholson, Thomas A. Püschel, Joanna Baker, Chris Venditti

## Abstract

A large brain relative to body mass is considered a distinguishing hominin trait. It has frequently been related to a suite of social, behavioral, technological, and other cognitive adaptations that differentiate humans from other species. The processes underlying large brain size evolution have therefore been a subject of rigorous scientific debate. Many hypotheses have been proposed to explain how climate and environment drive the selection of larger brain sizes, but monotonic influences of climate-environmental selective pressures are often assumed and rarely have between- and within-species effects been considered. Here, we apply Bayesian phylogenetic comparative techniques to the hominin fossil record to test the effect of climatic and environmental pressures (C-E) on brain size evolution, whilst simultaneously accounting for body mass and chronological age. We find that colder and more variable temperatures have a positive within-species effect on brain size evolution, likely related to biological adaptations to mitigate against hypothermia. However, in Homo, the strength of this effect diminishes over time suggesting that in later species (*Homo sapiens* and *Homo neanderthalensis*) brain sizes were less affected by C-E conditions.

## Introduction

Relative brain size is a particularly important trait as it is often used as a proxy for cognitive abilities^1–4^. It is widely reported that over the last few million years, relative brain size in hominins has increased, culminating in the iconic large brains of our own species^5^. However, relative brain size increase across hominin evolution arose from gradual change within individual species over time^6^. Hence, we must take a different approach from much previous research which only seeks patterns across species to truly understand the ecological drivers of brain size increase in hominins^4,7,8^.

Climatic and environmental (C-E) pressures have long been assumed to have played a crucial role on human encephalisation^3,4,9–14^. Consequently, multiple hypotheses have been proposed to explain the role of C-E variables (e.g., precipitation, temperature, vegetation) on hominin brain size evolution^3^. However, these hypotheses have been traditionally framed in ambiguous terms, leaving it unclear how they could be tested and with which data. More recently, Will et al.^3^ explicitly outlined hypotheses and associated expectations of how cranial capacity could be predicted from suites of so-called bioclimatic variables summarizing temperature and precipitation as well as a variable describing vegetation (henceforth net primary productivity, NPP). Briefly: The environmental stress hypothesis suggests that resource-deficient environments may induce stress-related brain size increases^3,15^ whereas the contrary environmental constraints hypothesis suggests that resource-rich environments are more likely to support an expensive, larger brain^3^. The environmental stress and environmental constraints hypotheses specifically predict opposing effects of temperature, precipitation and NPP on brain size. The environmental variability hypothesis predicts that increased cognitive abilities at short-time scales (or adaptive flexibility at longer timescales) are required to tolerate fluctuating environments^12^ whereas the environmental consistency hypothesis argues that climatic and environmental stability are more suited to maintaining large and metabolically costly brains^3,8^. The environmental consistency and environmental variability hypotheses make opposing predictions for variation in rainfall, temperature and NPP. All four hypotheses clearly outline the importance of either low/fluctuating resources or high/stable resources on varying timescales and make explicit predictions based on C-E data.

Whilst different studies find support for different hypotheses across the hominin radiation^3,4,8,15^, the data underlying the expectations of all hypotheses are not independent from one another (e.g.,^16^; Fig. S1), preventing clear conclusions from being drawn. Although bioclimatic variables and NPP are commonly used in studies of past environments and ecologies of extinct species, it is not possible to separate the effects of certain aspects owing to high levels of collinearity^17^. For instance, recent work has demonstrated that temperature, NPP, and precipitation all have similar impacts on body mass^3^. In short, the data underlying the predictions of current hypotheses seeking to explain the environmental drivers of hominin encephalisation are inextricably linked. It is necessary to understand this further before it is even possible to derive new, unbiased, and unambiguous environmental hypotheses of hominin brain size evolution.

Here, we take a unified approach to resolve outstanding questions about the influence of the climate and environment on hominin encephalisation. We reconstruct C-E data describing the quantity and variation of NPP, rainfall and temperature (including 17 bioclimatic variables; see SI and methods for full list of variables) for the given age and coordinates for each of 284 hominin specimens with cranial capacity using a climatic emulator spanning the last 5 million years of hominin evolution (see methods). This allowed us to specifically associate every analysed specimen with the C-E conditions of where they lived. To avoid collinearity among our predictors and reduce the dimensionality of our dataset we generate independent variables using principal component analysis (PCA). We then use our new orthogonal axes of C-E variation to formulate new hypotheses that can be tested in a Bayesian phylogenetic generalised linear mixed modelling (PGLMM) framework. To date, previous tests of the C-E drivers of hominin encephalisation have failed to 1) account for shared ancestry, 2) incorporate alternative phylogenies (^6^; Fig. 1B), 3) allow slopes to vary both between and within species, and 4) simultaneously account for within- and between-species effects of body mass and chronological age^6^. Our approach combines a reformulation of the C-E hypotheses and bioclimate proposed to drive hominin brain size evolution. We provide a robust novel perspective on encephalisation in hominins and the C-E conditions in which this occurred, whilst accounting for numerous sources of uncertainty (see methods).

**Figure 1.**
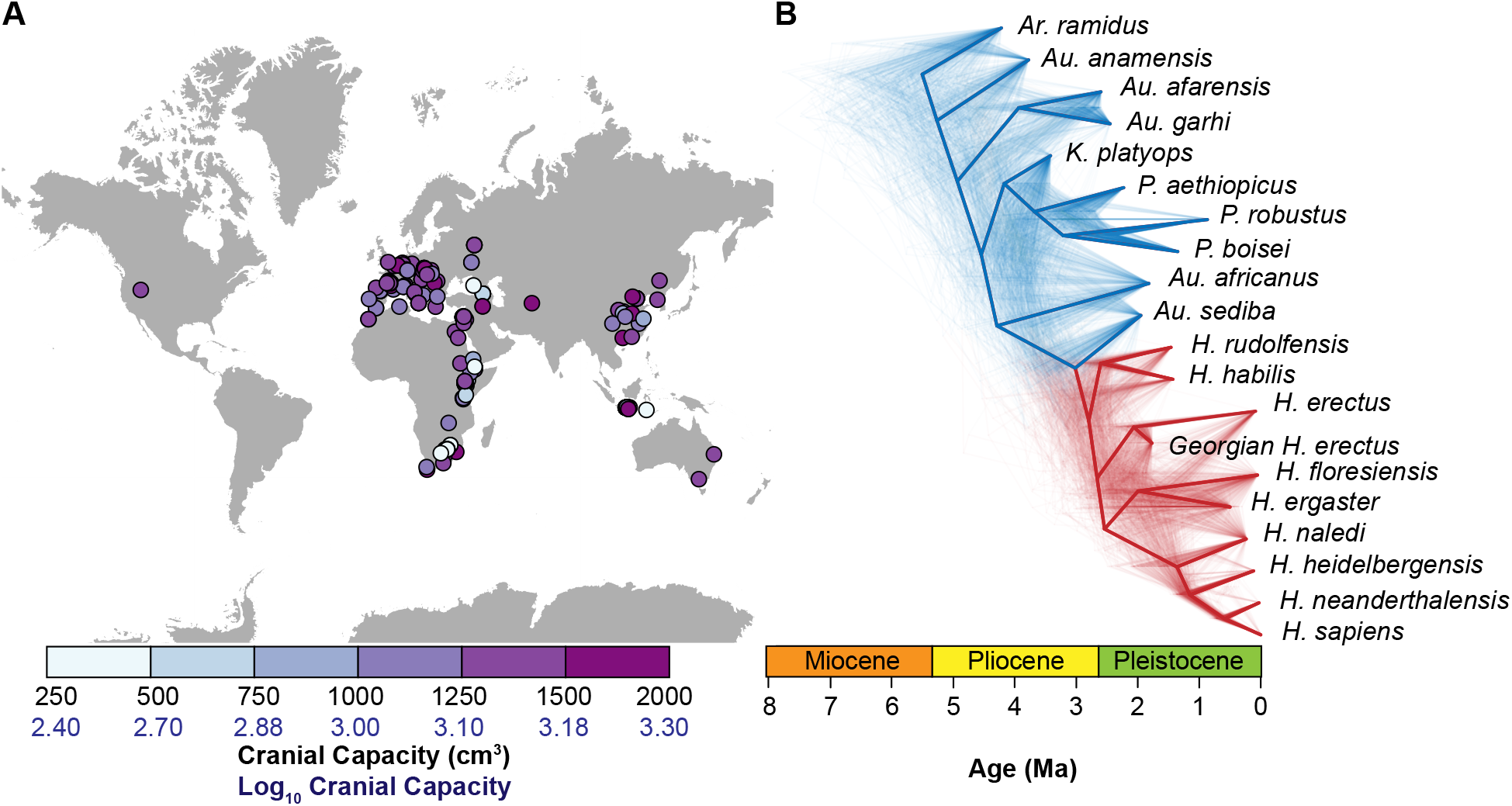
(A) Map of cranial capacity specimens used in this study and (B) Phylogenetic trees of the hominin radiation over the last 5 Ma. Thin lines represent a random sample (500) of alternative topologies, whilst the thicker topology corresponds to the hominin maximum credibility phylogenetic tree (MCC). Homo is colored in red, while the rest of the hominin genera are in blue.

## Results

### Climate principal component analysis

We reconstructed monthly mean temperature, precipitation and NPP for the last 5 Ma of hominin evolution (at 1-kyr intervals with a 0.5° resolution, see Methods) using the PALEO-PGEM climatic emulator^18^. From precipitation and temperature data, so-called “bioclimatic variables” - which describe the annual distribution and variation of rainfall, temperature and are independent of calendar months (for example, Bio-5 represents the temperature of the warmest month, rather than the temperature of August) and are widely used in ecological studies - were calculated following the ANUCLIM classification^19^. Additionally, to study long-term variance, we calculated the coefficient of variation for annual mean temperature, precipitation and NPP using a window of 23-kyrs to represent precession-scale variability (see SI for a complete list of these C-E variables). We then used point sampling procedures to extract these variables for each one of the hominins under study. We refer to all our data including the bioclimatic variables, NPP, and coefficients of variation as climatic-environmental (C-E) variables collectively.

We conducted PCA on a dataset comprising the C-E variables for 284 hominin specimens with cranial capacities (see methods). This process was repeated over a sample of 1,000 datasets, which allowed us to incorporate uncertainties related to body mass estimates, temporal range, and taxonomic assignment (^6^; see methods). Our first axis, PC1, explaining 36% of the variation in the data (Fig. 2A), is predominantly associated with temperature and can broadly be interpreted as a spectrum: low values describing warm environments with reduced temperature variability and range but with precipitation variability at both seasonal and long-term scales; up to high values, describing colder environments with greater temperature variability and range but with both annual and long-term precipitation stability (Fig. 2B). The second axis, PC2, accounts for ∼ 27% of the variance and is weighted more heavily towards precipitation: higher PC2 values indicate greater precipitation and NPP whereas lower values indicate drier environments.

**Figure 2.**
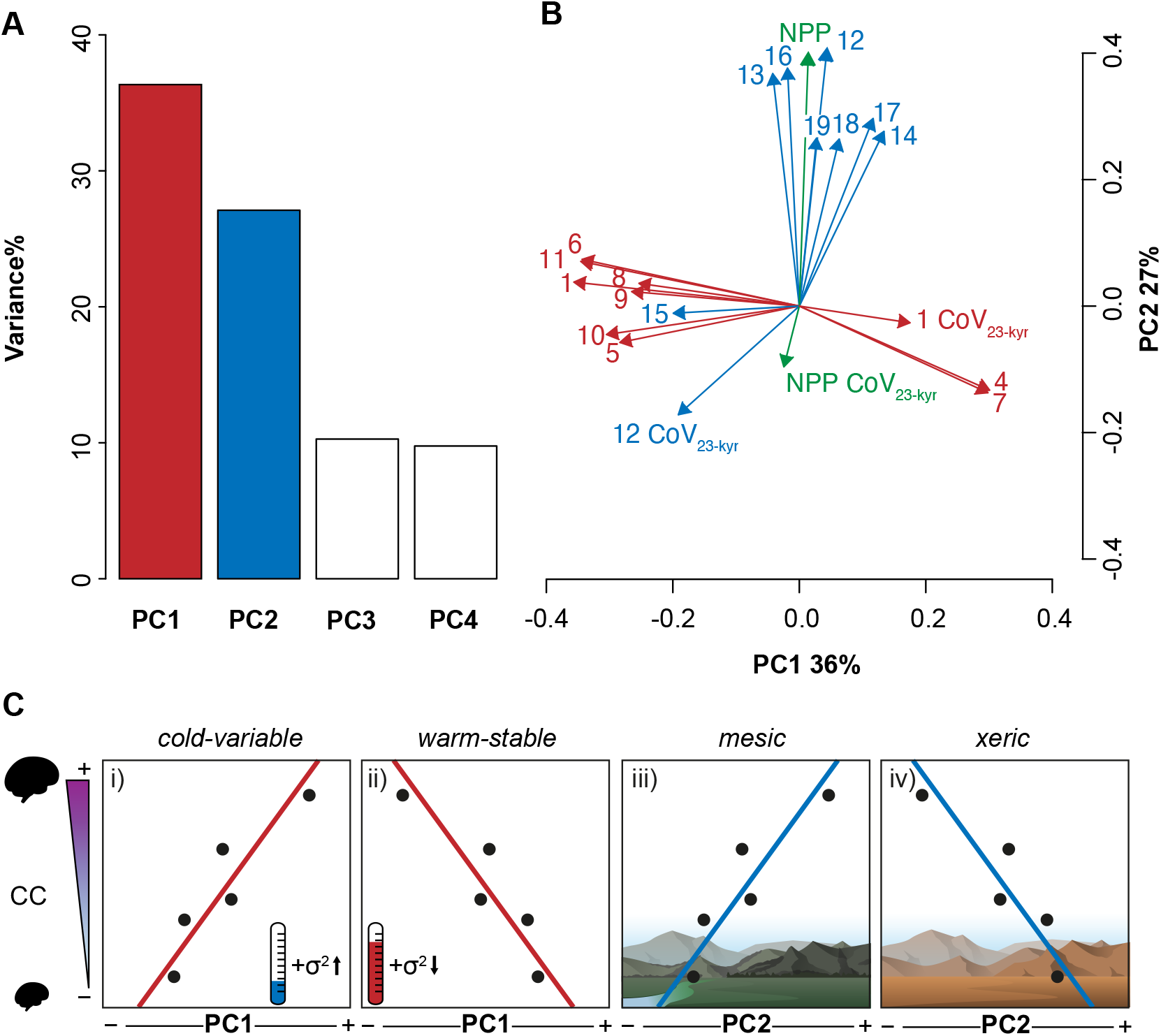
(A) Variance of the data explained by PCs. (B) Loadings of the bioclimatic variables on PC1 and PC2. Numbers relate to their respective bioclimate variables (see Tab. S1 for full list of variables). All values represent the mean over 1000 datasets. (C) Conceptual illustration of our new hypotheses for C-E influencers of brain size: i) the cold-variable hypothesis (where colder or more variable temperatures influence encephalisation), ii) the warm-stable hypothesis (where warmer or more stable temperatures influence encephalisation), iii) the mesic hypothesis (where wetter and more productive environments influence encephalisation), and iv) the xeric hypothesis (where drier and less productive environments influence encephalisation).

Seasonal variation is unimportant for PC2. Only these two first PCs were used in the subsequent analysis based on a parametric bootstrap method that was applied to determine the number of PCs to be kept^20^.

### The Climatic-Environmental drivers of hominin encephalisation

To test the expectations of the hypotheses we outline above, we conduct a phylogenetic generalised linear mixed model (PGLMM) testing the effect of each of our two PC variables on cranial capacity and whether this has been different within individual species. Our PGLMM approach enables us to account for body mass and chronological age whilst also allowing us to understand whether patterns differ both across and within individual species, taking into consideration phylogenetic relatedness. We find a significant positive relationship between cranial capacity within-species PC1 (Fig. 3), whilst there is no significant effect of PC2 on cranial capacity. Neither PC1 nor PC2 predicts brain size increase across species. In all models studied, we find a high phylogenetic signal (h2 = 0.88; see methods) and an overall good fit (R2marginal = 0.66; R2conditional = 0.95; see methods).

**Figure 3.**
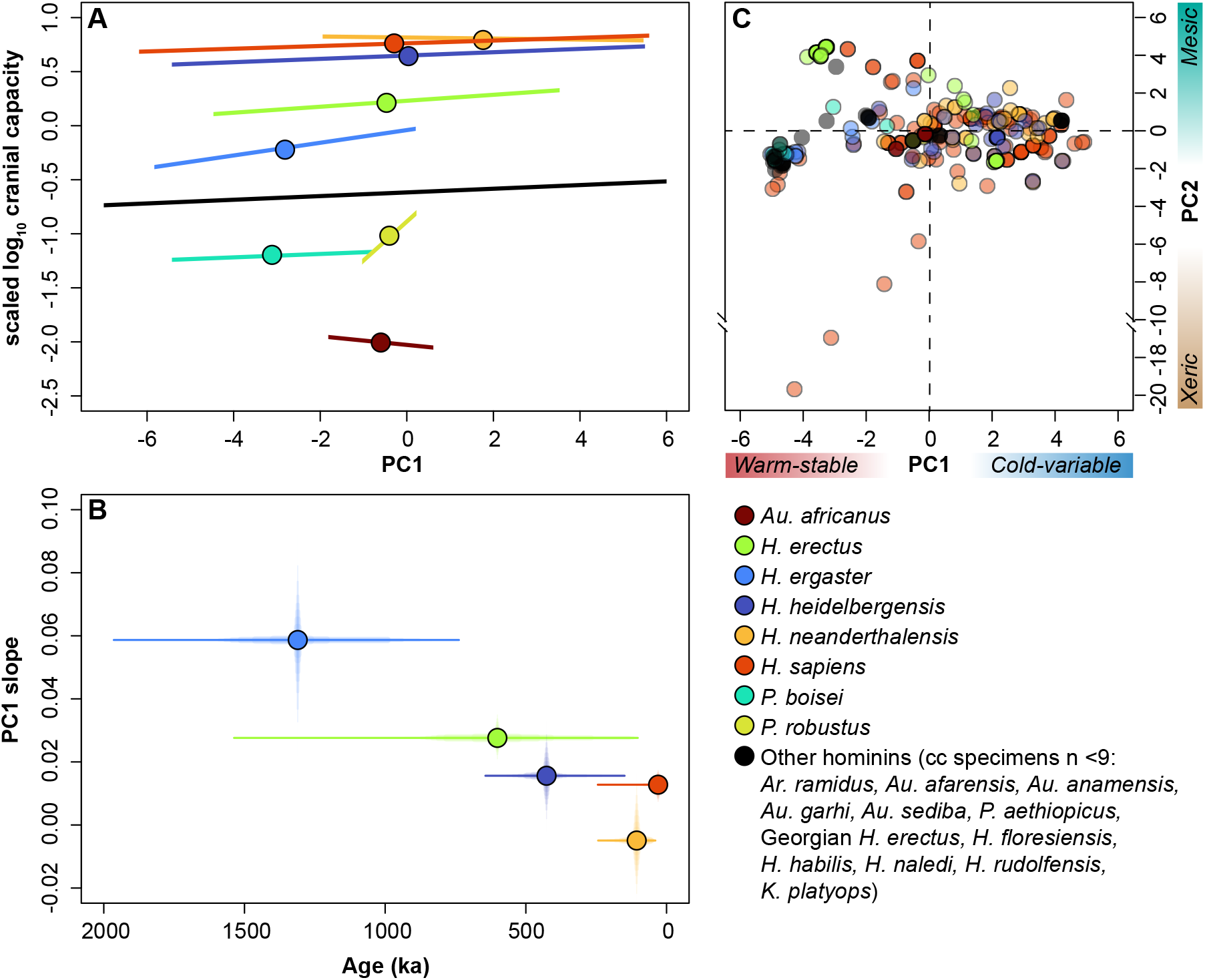
Fig. 3. (A) Within-species slopes (color lines) and overall, within-species slope (black line) for Scaled log10 cranial capacity vs. PC1. (B) Mean within-species slope and error (vertical lines) and species temporal durations (horizontal lines) through time for Homo species. Only species with >9 cc specimens are displayed. (C) PCA biplot of hominin cranial capacity specimens. Points represent the mean PC value over 1,000 datasets. *H. naledi* and *H. floresiensis* are not plotted on B as these do not have sufficient known temporal spans nor geographic information to accurately estimate their slopes.

## Discussion

The influence of predictor variables on hominin cranial capacity can invoke different modes of evolution (Fig. S2). Age and body mass have recently been studied and found that whereas body mass has a between-species effect, species age has a variable within-species. In other words, brain size increases occur at speciation events coeval to body mass increases and variably throughout the duration of a species (Fig S3: e,j and o). Here, we show that PC1 mostly has a positive effect on cranial capacity but there is a significant relationship within species (Fig. 3A).

This reveals a generally consistent trend supporting the predictions of the cold-variable hypothesis: within individual species, colder and/or more variable environments drive brain size increase, owing to stress. Furthermore, as between-species PC1 did not predict relative cranial capacity, we show that step changes in species average PC1 values do not influence brain size. Instead, it is the colder temperatures experienced in a species own range that enforce intraspecific increases in relative brain size. Variable within-species slopes for PC1 show that there is no monotonic influence of PC1 on relative brain size across the whole phylogeny: different species are affected differently by cold stress. Indeed, inversely, *Au. africanus* and *H. neanderthalensis* have seemingly negative slopes, suggesting warmer temperatures influence relative brain size in these species (Fig. 3A). Like previous findings^6^, between-species body mass and within-species age (with variable random slopes) also remain significant predictors of cranial capacity in hominins in our analysis. Another pertinent observation is that age remains significant, even after accounting for body mass and C-E conditions, implying that other processes throughout the duration of a species’ existence also influence relative cranial capacity increases. In other words, C-E conditions are not the only means to increase hominin cranial capacities.

Unknown factors not studied here have additional influence and future investigation will be required to elucidate these. Together, these reveal a pattern whereby increases of cranial capacity aligned with body mass increases occur speciation events (Fig. S2: j); subsequent within-species encephalisation occurred due to colder conditions (Fig. 3A) and unknown factors consistent with age.

But how do low/variable temperatures induce brain size increases? Previous attempts to directly associate colder temperatures and/or temperature variability with encephalisation in primates and hominins have focused on biological^8,21–23^, social^23^ and techno-cultural solutions to prevent hypothermia. It is tempting to link this to Bergmann’s rule, where larger organisms are typically related to higher latitudes. Indeed, temperature correlates with latitude^24^. However, Bergmann’s rule is related to total body mass, which was explicitly accounted for in our analysis. Furthermore, we ran an independent test for relative latitude (i.e., degrees distance from the equator) and, contrary to other studies^8^, it was not a significant predictor of hominin brain size within- or between-species (Tab. S1). Moreover, most attempts to link temperature to encephalisation have also assumed that temperature controls “resources”, where colder/variable environments are more resource-scarce/variable, thus creating selective pressure for enhanced cognitive capabilities ^3^. However, we found a low correlation between temperature indices and NPP (i.e., a proxy for productive environments; Fig. S1); NPP neither loads highly on PC1, nor does PC2 (with a high NPP loading) predict hominin brain size. Instead, NPP is highly correlated with precipitation indices (Fig. S1). Our results suggest the direct effects of ambient absolute temperature/variability influence brain size evolution.

One potential mechanistic explanation for the observed relation between relative brain size and PC1 values representing temperature can be advanced by the so-called “radiator hypothesis” ^21^. This hypothesis stipulates that a network of emissary veins evolved to cool the brain under conditions of hyperthermia, and diploic veins that provide a mechanism for cooling have been associated with larger brains. Colder environments typically have greater seasonal temperature oscillations; fluctuations between hyper- and hypothermic conditions within a species’ environment may have provided a mechanism for encephalisation by demanding an enhanced diploic vein system for effective thermoregulation^8^. Whilst the radiator hypothesis is heavily linked to the evolution of bipedal posture, climate may have influenced its evolution by providing a stimulus for more efficient thermoregulation^21^, particularly for those species with negative PC1 slopes.

But the radiator hypothesis primarily describes the prevention of hyperthermic conditions ^21^ – what about colder temperatures? Recent studies have shown that vertebrates typically have larger brains in colder conditions, even when accounting for body mass^25^. Additionally, large brains in Asian colobine monkeys have been shown to have evolved during colder events and resultant cold adaptations may have influenced social organisation^23^. In *H. sapiens* larger brain sizes seemingly occurred with globally cooler periods^15^. If colder events influence brain size, we must establish the limits at which temperature stress could occur. Past studies have provided minimum sustainable temperatures for Homo erectus of 11.67°C and 6.27°C^26,27^. With the exemption of tropical latitudes, this is well within the temperature variation experienced during cold events (Fig. S2). Assuming, based on smaller body masses, this value was higher in earlier species, cold may have been a long-term stress throughout hominin evolution. Thus, this explanation would work within-species, as it would take effect within any species’ own geographic range where colder areas are more likely to induce species-specific stress. We also show that global mean annual temperature did not predict brain size (Tab S2), reinforcing our suggestion for an influence of temperature in local environments. Yet, further information on the minimum sustainable temperatures across the hominin radiation is required to truly elucidate the mechanisms of cold stress-induced encephalisation. Previous studies^3^ found that, whereas body mass evolved as a phenotypic adaptation to cope with short-lived low temperatures, brain size was related to variations in precipitation and NPP. Our results however, which explicitly accounted for the correlation between cranial capacity and body mass, rather show that relative brain size also evolved to cope with stress induced by short-lived low temperatures, but this effect varied between species. The exact processes of how colder-variable temperatures influence brain size is not clear and likely complicated. Owing to the high correlation of temperature variables, a simple “cold causes larger brains” does not sufficiently explain the within-species variance of the effect and its influence on even low-latitude hominin species.

One stark observation that we make is that, within later species of Homo, PC1 is having a lesser impact on encephalisation. There is a significant correlation (r = 0.95, *p* = 0.02) between median age (ka) and the median slope estimation. The median slope estimates decline from ∼0.06 in *Homo ergaster* to ∼0.01 in *Homo sapiens*, and fall to ∼-0.01 in *Homo neanderthalensis*. However, *H. sapiens* and *H. neanderthalensis* have a more rapid encephalisation throughout their temporal duration when compared to earlier species^6^. One potential climatic explanation for this encephalisation is that increased temperature variability related to greater glacial-interglacial amplitudes, following the Mid-Bruhnes event (∼460 ka) could have enhanced climatic pressures on later *Homo* species^28^. Indeed, broad narratives of global Quaternary climate change include a transition from orbitally-paced 41-kyr “obliquity” to 100-kyr “eccentricity” interglacial-glacial cycles between 1.2-0.7 Ma^29–32^, followed by increased amplitude and duration of glacials following the Mid-Bruhnes event^28^. These changes are seemingly coeval to increases in maximum hominin brain sizes. We therefore conducted further PGLMM analyses using the moving Coefficient of Variation on orbital cycles (precession [long]: CoV23-kyr; obliquity: CoV41-kyr; eccentricity: CoV100-kyr) for Bio-1 (Mean Annual Temperature), Bio-12 (Annual Precipitation) and NPP (Net Primary Productivity) (methods). Neither the CoV of precipitation and NPP on precession (∼23-kyrs), obliquity (∼41-kyrs) and eccentricity (∼100-kyrs) timescales (Fig. S3) predicted relative brain size (Tab. S3). On the other hand, Bio-1 CoV100-kyr was found to be significant. Additional testing, however, found a strong correlation with PC1 (0.75) and these effects cancelled each other when modelled in tandem (Tab. S3). The effects of PC1 and Bio-1 CoV100-kyr are therefore indistinguishable, and we cannot relate this Mid-Late Pleistocene rapid encephalisation to enhanced glacial-interglacial climatic variability in temperature, precipitation or NPP. It is likely that alternative factors are influencing encephalisation within these later hominin species.

One consideration is that these later Homo species have large geographic ranges^33^. Preceding species, such as *H. ergaster* (southern and eastern Africa) and *H. erectus s*.*s*. (south-eastern and eastern Asia) appear to have more restricted regional distributions. Conversely, depending on controversial species assignments, *H. heidelbergensis* may have been distributed across Africa, Europe, and Asia. Likewise, Neanderthal specimens were recovered in Europe and south-western Asia and *H. sapiens* specimens were on every continent (except Antarctica) by the end of the Pleistocene (11.7 ka). Both Neanderthals and *H. sapiens* plot more widely on PC1 and PC2 (Fig. 3C) and reflect greater C-E variation. Such a range of C-E conditions may have abated the importance of any one climatic metric, suggesting that the multiplicity of climates and environments may have contributed to Mid-Late Pleistocene encephalisation. Another consideration is that techno-cultural (i.e., fire, clothing, shelter-building, other survival/subsistence strategies^26,34–36^) and socio-behavioral innovations (i.e., language, symbolism, group-size changes^37^) may have provided mechanisms (e.g., niche construction) that abated the C-E effects influencing encephalisation, or perhaps were alternative inducers of encephalisation.Understanding this trend should be a target for future research, where the effect of biogeographic and C-E range throughout a phylogeny can be tested against brain size.

As shown here in our work, countless climatic hypotheses can be envisioned but only a more limited number of them can be statistically assessed using the current evidence from the fossil record^3,4,8^. From our reformulation of C-E encephalisation hypotheses, our analysis supports a scenario in which absolute temperature and temperature variability within a species’ own geographic range influenced encephalisation. Importantly, the differential effects of climate at the species level mitigate against broad narratives of climatic impacts on hominin evolution, such as previously proposed hypotheses^4,11–13,38,39^. Our results show a diminishing impact of climate within the later species of large-bodied Homo, including *H. sapiens*, as total geographic ranges expanded and technological/behavioral innovations increased^34^. We, therefore, emphasise against a simple determinist view of climatically driven cognitive advances for these species. Our findings contrast recent studies, and we highlight the necessity of studying climate impacts on cranial capacity whilst accounting for between- and within-species effects, phylogenetic relatedness, body mass and chronological age within a single inclusive framework.

## Materials and Methods

### Hominin sample and phylogenies

We used the fossil dataset of hominin specimens with cranial capacity from Püschel et al.^6^ (supp. data 1). However, we excluded *S. tchadensis*, as the PALEO-PGEM climate emulator does not extend back to its reputed age of ∼7.2 Ma^40^. This resulted in a sample size of 284 hominin specimens ranging from ∼ 4.4 Ma to the end of the Pleistocene. We associated every specimen with cranial capacity to a body mass value and chronometric age using well-defined criteria (see^6^ for details), which allowed us to consider uncertainties related to both body mass estimates, taxonomic assignment, and temporal range. This process was repeated 1,000 times resulting in 1,000 unique datasets. We also provided geographical coordinates, recorded as longitude and latitude ^6^, for each one of the hominin specimens under study.

A sample of 1,000 phylogenetic trees was obtained from Püschel et al.^6^ (supp. data 2). These trees were randomly sampled from a posterior distribution of phylogenies obtained using a ‘combined-evidence’ Bayesian phylogenetic reconstruction of hominin species using stratigraphic, molecular, and morphological data. These phylogenetic trees were used in our subsequent analyses (see PGLMM section).

### Climate-Environmental Data

We reconstructed environmental data for the last 5 Ma starting with climatic variables generated using the PALEO-PGEM climatic emulator. PALEO-PGEM applies a Gaussian process emulation to singular value decomposition of ensemble runs of the intermediate complexity atmosphere-ocean general circulation model PLASIM-GENIE21,42. Boundary condition forcings include atmospheric CO2 (after 800 ka^41^; before 800 ka^42^), sea-level (as a proxy for ice-sheet)^42^ and orbit (obliquity and precession-modulated eccentricity)^43^. Orography is fixed. The native-resolution (5°) emulations have previously been validated against of four discrete periods: the mid-Holocene, the Last Glacial Maximum, the Last Interglacial, and the mid-Pliocene warm period^18,44^. The PALEO-PGEM climate emulator was run via R (R version 4.1.2^45^) on the default gaussian-exponential setting, following the same procedures as Raia et al.^44^. In this setting, the PALEO-PGEM uses a power exponential covariance function, the mean GP prediction and ten principal components.

Spatial fields of temperature, precipitation and annual Net Primary Productivity are then emulated at 1,000-year intervals, driven by time series of scalar boundary condition forcing, and assuming the climate is in quasi-equilibrium^18,44^. However, unlike Raia et al.^44^ – who ascertained minimum seasonal temperature, maximum seasonal temperature, minimum seasonal precipitation, and maximum seasonal precipitation – we calculated monthly temperature and precipitation values.

PALEO-PGEM was run ten times for each monthly temperature and precipitation variable and a mean was taken for the analysis. Following this, outputs were converted to 1-kyr rasters of 5° resolution using a Thin Plate Spline interpolation (e.g.,^46^). These were then downscaled to 0.5° resolution and bias-corrected using the respective climate rasters averaged over the 1981-2010 period available from the CHELSA database48–50^47–49^. Downscaling of temperature, precipitation and NPP followed previously established procedures^18,44^, where an additive correction was used for temperature and a conditional hybrid (additive or multiplicative) correction was used for precipitation and NPP (see^18,44^ for full description) using the following formulae:

Additive correction:

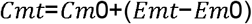

Multiplicative correction:

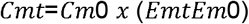

Where Emt is the pre-downscaled emulator output at time (t), Em0 is the pre-downscaled emulator at 0 ka, Cm0 is the climate variable at 0 ka by which downscaling should take place, and Cmt is the downscaled climate variable at t. For the hybrid procedure, an additive correction is employed if Cm0>Em0; whereas a multiplicative procedure is employed if Cm0<Em0. This downscaling procedure aims to minimize negative precipitation values in hyper-arid (deserts) areas and unrealistically high values in hyper-humid areas (tropical rainforests). However, where we identified instances of negative precipitation values in deserts, these were manually adjusted to 0. Raw outputs from the emulator are measured as Celsius and mm day^-1^ for monthly temperature and precipitation, respectively, and kg C m2 s^-1^ (kilograms of carbon per square meter per second) for annual NPP. These were transformed to monthly Kelvin (temperature), monthly absolute mm (which in this case equates to mm month-1) for precipitation, and g C m^2^ yr^-1^ (grams of carbon per square meter per year) for NPP.

Following downscaling, we calculated 17 of the 19 bioclimatic variables defined in the ANUCLIM classification ^19^. Two bioclimatic variables, Bio-2 (Mean Diurnal Range) and Bio-3 (isothermality) ^19^, were not calculated as these require monthly minimum and maximum temperatures and presently PALEO-PGEM can only predict monthly mean temperatures. Additionally, for bioclimatic variables where monthly minimum or maximum temperature are given (Bio-5: maximum temperature of the warmest month; Bio-6: minimum temperature of the coldest month; Bio-7: annual temperature range (Bio-5 – Bio-6), these are instead mean temperatures of the hottest/coldest month. Next, we calculated the moving Coefficient of Variation (CoV = standard deviation/mean) for annual mean temperature and precipitation, as well as NPP using a window of 23-kyrs to represent long precession-scale variability. This resulted in a total of 21 variables that were used to characterize past environments.

We then used coordinate and age data to sample the appropriate location on each 1-kyr map of each of the 21 environmental variables, which allowed us to link every specimen from the 1,000 unique datasets to the conditions of where they lived. This procedure resulted in 1,000 datasets which included each specimen with a chronometric age (constrained by the specimen age range) with associated cranial capacity, body mass and environmental data (supp. data 3 and supp. data 4). Cranial capacity, body mass and all environmental variables (excluding CoV) were log10 transformed. Finally, all data was standardized (i.e., centered and scaled) prior to analysis.

### Environmental PCA

We carried out a PCA using all the environmental variables to reduce the dimensionality of our dataset and avoid predictor collinearity using the R (version 4.1.2) statistical software package ^45^. This PCA was performed using the correlation matrix of the 17 bioclimatic variables, NPP and CoV23-kyr for Bio-1, Bio-12 and NPP (Fig. S1), which allowed us to generate new orthogonal axes of variation (PCs) that were subsequently used as independent environmental variables in our analyses. To determine the number of PCs to be retained in our further analytical steps, we applied a full parametric bootstrap method recommended for standardized variables ^20^. All these procedures were repeated 1,000 times using the 1,000 datasets previously mentioned.

### Bayesian phylogenetic generalised linear mixed modelling

Each of the 1,000 datasets were analysed using Bayesian phylogenetic generalised linear mixed models (PGLMMs) to test the relationship between cranial capacity and the within- and between-species effect of body mass, chronological age, and environment (i.e., PC1 and PC2). A PGLMM is like a conventional phylogenetic generalised least squares (PGLS) method but goes beyond merely estimating the variance of the phylogenetic effect: It also incorporates an additional residual error term which allows for the inclusion of other factors such as intraspecific variance, environmental effects, measurement error, and more. PGLMM incorporates shared ancestry information by including a phylogenetic random effect that is assumed to follow a normal distribution with a variance that considers correlation among phylogenetic effects based on a phylogenetic variance-covariance (or correlation) matrix. In our case, we computed these matrices using the 1,000 hominin phylogenies previously mentioned.

To assess the role of inter- and intra-specific variability, we used a technique known as within-group centering ^50^. This technique partitions the predictor variables into two different components: the group-level mean (i.e., the mean of that predictor for each species, resulting in the between-species differences) and the within-group variation (i.e., the difference of each subject from its within-group mean, resulting in the within-species differences). We also accounted for possible slope differences per species (i.e., a random slope model) by adding different random effects (i.e., species-specific random effects for the within-group variability of both body mass, time and environment represented by PC1 and PC2).

For each bioclimatic variable and PC, we conducted PGLMM analyses with a set random-effect structure (supplementary files). As our previous work has demonstrated that between-species body mass and within-species chronological age are significant predictors of hominin cranial capacities ^6^, all regressions included between- and within-species body mass, chronological age, and PCs as fixed effects (supplementary files). Within-species predictors were also included as random effects. In addition, we ran all bioclimatic variables following this structure (Tab. S1).

Results for other significant climate variables (Bio-1, -4, -6, -7 and -11), as well as the results of non-significant climate variables are included in the supplementary files (Tab. S1). All reported values from our modelling approach correspond to grand means (i.e., average of means) computed for the estimated effects obtained by the models ran using our 1,000 unique datasets (each using an alternative phylogenetic tree).

PGLMM analyses were conducted using the MCMCglmm v.2.33 R package ^51^. For the fixed effects, we used a diffuse, normally distributed, prior centered around zero (μ=0) with very large variance (σ2=10^8). Inverse-Gamma prior distributions with shape (α) and scale (β) parameters equal to 0.01 were applied for the random effects. All regressions were conducted using a total of 510,000 iterations, with a thinning interval of 250, and the first 10,000 iterations were removed as burn-in. Markov chain convergence and mixing were visually assessed by looking at the trace plots of each one of the fixed and random effects and we checked that effective sample sizes were > 1,000. Significance was assessed by the proportion of the posterior distribution crossing zero (Px). Where Px < 0.05, less than five percent of the posterior distribution overlaps with zero, and we consider a predictor to be different from zero.

## Acknowledgements

This work was supported by Leverhulme Research Leadership Award RL-2019-012

## Notes

**Competing Interest Statement:** The authors declare no conflicts of interest.

### Competing Interest Statement

The authors have declared no competing interest.

