## Supplementary file for "Climatic-environmental influences on hominin brain size over the last 5 million years"

### PGLMM analysis and results

PGLMM analysis was conducted using 17 bioclimatic variables, NPP, the coefficient of variation on a moving 23-kyr (long precession cycle) window (CoV_23-kyr_), as well as the first four PCs obatined from a PCA computing using these variables (Tab. S1). To account for the within- and between-species effects, as well as the effect of within- and between-species effects of body mass and chronometric age, the tests included the following structure:

Cranial capacity ~ species mean body mass: within species body mass: species mean age: within species age: species mean test: within species test.

In addition, a random effect term was included to account for phylogenetic relatedness, and the within-species slope variation for body mass, age and the test. Significant (pMCMC < 0.05) climatic variables included bio-1 (mean annual temperature), -4 (annual temperature seasonality), -6 (mean temperature of the coldest month), -7 (annual temperature range) and -11 (mean temperature of the coldest quarter), where lower mean temperatures or greater temperature variability predicted larger relative brain-size. (Phylogenetic signal and marginal and conditional R^2^ values are available in supplementary information). In all significant tests (pMCMC <0.05), between-species body mass and within-species age remained significant predictors of brain-size, providing additional support for previous findings and highlighting the importance of these factors. The significant climate variables are highly correlated to each other (Fig. S1), where lower temperatures are associated to greater annual temperature variability and range. In contrast to when analysed independently, when analysed together these variables cancel out the significance of each other due to variance inflation (1). Additionally, temperature variables are all highly colinear with relative latitude (2). We therefore run an independent phylogenetic regression for relative latitude. Yet, we find that latitude is not a significant predictor of relative brain-size (Tab. S3). This indicates that local environmental temperature variability within a species’ biogeographic range drives brain-size increase. This is in stark contrast to a recent study that found that low temperatures predict body mass, and that cranial capacity was predicted by NPP and long-term precipitation variability (3). Our method, which considers the collinearity between climate variables, between-species effect of body mass and within-species effect of chronology, and phylogenetic relatedness, found that only temperature and temperature variability is a significant predictor of hominin relative brain-size.

There is species-specific variation in slopes between temperature variables related to absolute temperatures (bio-1, -6 and -7) or temperature variability (bio-4 and -7). This suggests that 1) whilst there is high collinearity between temperature variables across the whole dataset, this trend is not as apparent within a species’ own geographic range; and 2) different temperature variables affect species differently and implies that different hypotheses explain brain size evolution in different species but are indistinguishable across the whole phylogeny.

| **Test variable** | **Description** | **Intercept** | **Between-species var slope** | **Between-species var pMCMC** | **Within-species var slope** | **Within-species pMCMC** | **R2 conditional** | **R2 marginal** | **h^2^** |
| --- | --- | --- | --- | --- | --- | --- | --- | --- | --- |
| **log10 Bio-1** | **Mean Annual Temperature** | **-0.60** | **-0.05** | **0.32** | **-0.05** | **0.01** | **0.95** | **0.66** | **0.88** |
| **log10 Bio-4** | **Temperature Seasonality** | **-0.63** | **0.03** | **0.36** | **0.04** | **0.03** | **0.95** | **0.66** | **0.88** |
| log 10 Bio-5 | Mean temperature of the warmest month | -0.65 | -0.01 | 0.38 | -0.03 | 0.08 | 0.95 | 0.65 | 0.88 |
| **log10 Bio-6** | **Mean temperature of the coldest month** | **-0.60** | **-0.05** | **0.31** | **-0.05** | **0.01** | **0.95** | **0.67** | **0.88** |
| **log10 Bio-7** | **Annual temperature range** | **-0.58** | **0.02** | **0.36** | **0.04** | **0.03** | **0.94** | **0.66** | **0.88** |
| log10 Bio-8 | Mean temperature of the wettest quarter | -0.57 | -0.08 | 0.28 | -0.04 | 0.06 | 0.95 | 0.68 | 0.88 |
| log10 Bio-9 | Mean temperature of the driest quarter | -0.66 | 0.00 | 0.37 | -0.02 | 0.08 | 0.95 | 0.65 | 0.88 |
| log10 Bio-10 | Mean temperature of the warmest quarter | -0.64 | -0.01 | 0.37 | -0.03 | 0.06 | 0.95 | 0.65 | 0.88 |
| **log10 Bio-11** | **Mean temperature of the coldest quarter** | **-0.60** | **-0.05** | **0.32** | **-0.05** | **0.01** | **0.95** | **0.67** | **0.88** |
| log10 Bio-12 | Annual precipitation | -0.57 | 0.00 | 0.24 | 0.00 | 0.44 | 0.95 | 0.66 | 0.88 |
| log10 Bio-13 | Precipitation of the wettest month | -0.49 | 0.00 | 0.18 | 0.01 | 0.34 | 0.95 | 0.69 | 0.87 |
| log10 Bio-14 | Precipitation of the driest month | -0.62 | 0.00 | 0.27 | -0.02 | 0.21 | 0.96 | 0.67 | 0.87 |
| log10 Bio-15 | Precipitation Seasonality | -0.42 | 0.00 | 0.12 | 0.02 | 0.13 | 0.96 | 0.74 | 0.86 |
| log10 Bio-16 | Precipitation of the wettest quarter | -0.50 | -0.19 | 0.18 | 0.01 | 0.36 | 0.95 | 0.68 | 0.87 |
| log10 Bio-17 | Precipitation of the driest quarter | -0.64 | 0.08 | 0.31 | -0.01 | 0.28 | 0.95 | 0.66 | 0.88 |
| log10 Bio-18 | Precipitation of the warmest quarter | -0.50 | -0.39 | 0.11 | 0.00 | 0.42 | 0.95 | 0.71 | 0.87 |
| log10 Bio-19 | Precipitation of the coldest quarter | -0.62 | 0.19 | 0.21 | -0.02 | 0.20 | 0.95 | 0.69 | 0.87 |
| log10 NPP | Net Primary Productivity | -0.45 | -0.21 | 0.19 | 0.00 | 0.39 | 0.94 | 0.68 | 0.87 |
| CoV bio-12 | 23 kyr moving coefficient of variation of annual precipitation | -0.40 | -0.20 | 0.10 | -0.02 | 0.14 | 0.94 | 0.72 | 0.85 |
| CoV NPP | 23 kyr moving coefficient of variation of NPP | -0.56 | -0.06 | 0.29 | 0.02 | 0.19 | 0.94 | 0.67 | 0.87 |
| CoV bio-1 | 23 kyr moving coefficient of variation of mean annual temperature | -0.61 | -0.05 | 0.31 | 0.02 | 0.17 | 0.94 | 0.65 | 0.87 |
| **log10 PC1** | **Temperature** | **-0.60** | **0.02** | **0.32** | **0.02** | **0.02** | **0.95** | **0.66** | **0.88** |
| log10 PC2 | Precipitation and productivity | -0.60 | -0.05 | 0.30 | 0.00 | 0.33 | 0.95 | 0.66 | 0.88 |
| log10 PC3 | Minimum precipitation | -7.90 | -0.02 | 0.20 | 0.01 | 0.19 | 0.94 | 0.70 | 0.86 |
| log10 PC4 | Noise | -7.66 | 0.00 | 0.16 | -0.01 | 0.16 | 0.95 | 0.71 | 0.86 |

*Tab. S1. Table of results of PGLMM analysis. All tests included between- and within-species body mass, chronological age and the tested climate variable as fixed effects. Random effects included phylogeny and the species-specific within-species body mass, chronological age and the studied climate variable. The regression structure follows: Cranial capacity ~ species mean body mass: within species body mass: species mean age: within species age: species mean test: within species test.*

| **Test Variable** | **Between-species BM slope** | **Between-species BM pMCMC** | **Within-species BM slope** | **Within-species BM pMCMC** | **Between-species age slope** | **Between-species age pMCMC** | **Within-species age slope** | **Within-species age pMCMC** |
| --- | --- | --- | --- | --- | --- | --- | --- | --- |
| log10 Bio-1 | 0.52 | **0.005** | 0.04 | 0.10 | 0.00 | 0.07 | 0.00 | **<0.00** |
| log10 Bio-4 | 0.51 | **0.005** | 0.04 | 0.08 | 0.00 | 0.07 | 0.00 | **<0.00** |
| log10 Bio-6 | 0.51 | **0.005** | 0.04 | 0.10 | 0.00 | 0.07 | 0.00 | **<0.00** |
| log10 Bio-7 | 0.51 | **0.005** | 0.04 | 0.08 | 0.00 | 0.07 | 0.00 | **<0.00** |
| log10 Bio-11 | 0.51 | **0.005** | 0.04 | 0.10 | 0.00 | 0.07 | 0.00 | **<0.00** |
| PC1 | 0.52 | **0.005** | 0.04 | 0.10 | 0.00 | 0.07 | 0.00 | **<0.00** |
| CoV Bio-1 _100-kyr_ | 0.53 | **0.005** | 0.04 | 0.11 | 0.00 | 0.08 | 0.00 | **<0.00** |

*Tab. S2. Slope and pMCMC values for the within- and between-species effects of body mass (BM) and chronometric age (age) for the significant tests in Tab. S1 and Tab. S3.*

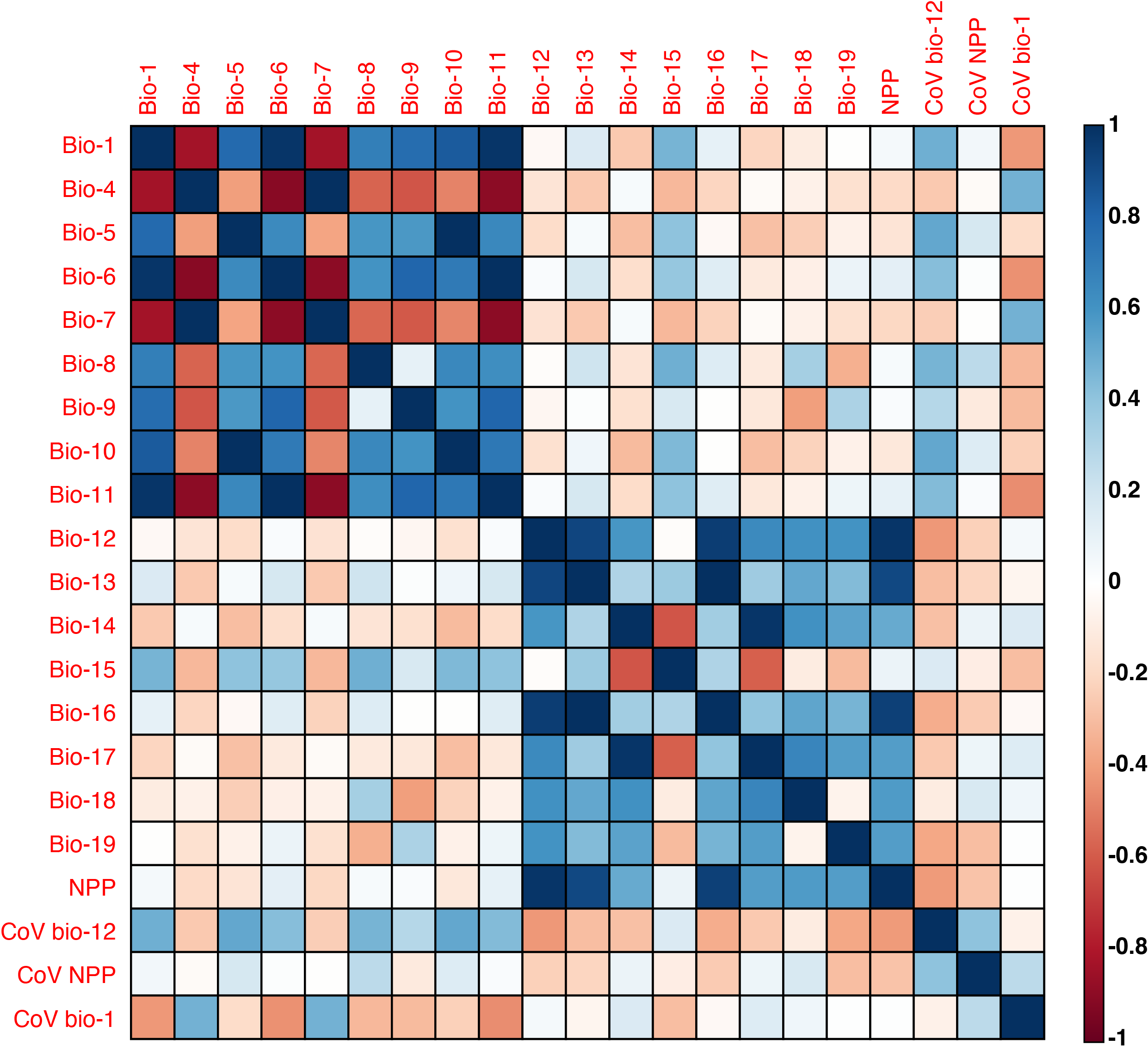

*Fig. S1. Correlation matrix of mean correlation coefficients of bioclimate variables, NPP and CoVs over our 1000 datasets.*

### Cold stress and distribution of global distribution of minimum temperatures

Much archaeological work, especially for the Lower-Middle Palaeolithic, has attempted to establish the climatic tolerances of humans in the context of fluctuating interglacial-glacial conditions. In north-western Europe, for example, where the temperature variance between glacial and interglacial conditions is large, discussions have focussed on survival strategies to survive extreme cold and prevent hypothermia (e.g., (4, 5)). In modern humans, hypothermic conditions can be onset by temperatures less than 10^o^C. Minimum sustainable temperatures (MSTs) for past humans have been used to estimate at which temperature additional clothing would be required to prevent hypothermic conditions. Aiello and Wheeler (6) estimated MSTs of 10.5-4.9^o^C for *Homo sapiens* and 11.6-6.2^o^C for *Homo erectus*. Across areas of Eurasia, modelled winter temperatures throughout the Lower-Middle Pleistocene frequently fall below these temperatures. For example, at Harbin, the locality of the so-called “Dragon Man” specimen HBSM2018-000018(A), the lowest annual minimum temperature over 1909 to 1990 averaged -26.7^o^C (data from KNMI climate explorer, Harbin station; https://climexp.knmi.nl/getmin.cgi?WMO=50953) and the average estimate for Bio-6 (temperature of the coldest month) from PALEO-PGEM over its dating range was -21.5^o^C. Various techno-cultural (e.g., fire and clothing) and subsistence strategies (e.g., long-range migration) have been proposed to overcome such temperature extremes (4).

As PC1 has a positive influence on brain size, this suggests a causal mechanism between colder or more variable temperatures and increased brain sizes. This positive relationship, however, is also present in low-latitude species (opposed to the Neanderthals whose range spread across Europe, the Middle East and northern Asia), mitigating against a simple paradigm that larger brains could only evolve at high latitudes. This therefore raises the question about how often lower-latitude hominins may have experienced hypothermic temperatures. It should be noted that hypothermia can onset within hours, and the 1000-year time-averaged minimum temperatures from the PALEO-PGEM (7, 8) emulator is unlikely to capture extreme minimum temperature “cold-snap” events. We must therefore use finer resolution climate data to understand how minimum temperatures are distributed globally. We searched the KNMI climate explorer for minimum daily temperature observations since 1950. We collated the monthly minimum temperatures over this period and created a raster map displaying the lowest recorded temperature over the entire period (Fig. S2). Assuming that cold stress drives selection for brain size in hominins, we need only find that temperatures can fall below minimum sustainable temperature. As can be observed, may places – even stereotypically “hot” environments (e.g., tropical rainforests and deserts) – can experience temperatures that fall within or below the MST of *H. erectus*. Indeed, even in the Saharo-Arabian desert belt, winter night-time temperatures can fall below freezing (9). Minimum temperatures over 11.6^o^C are mostly centred within the tropics (Fig S2).

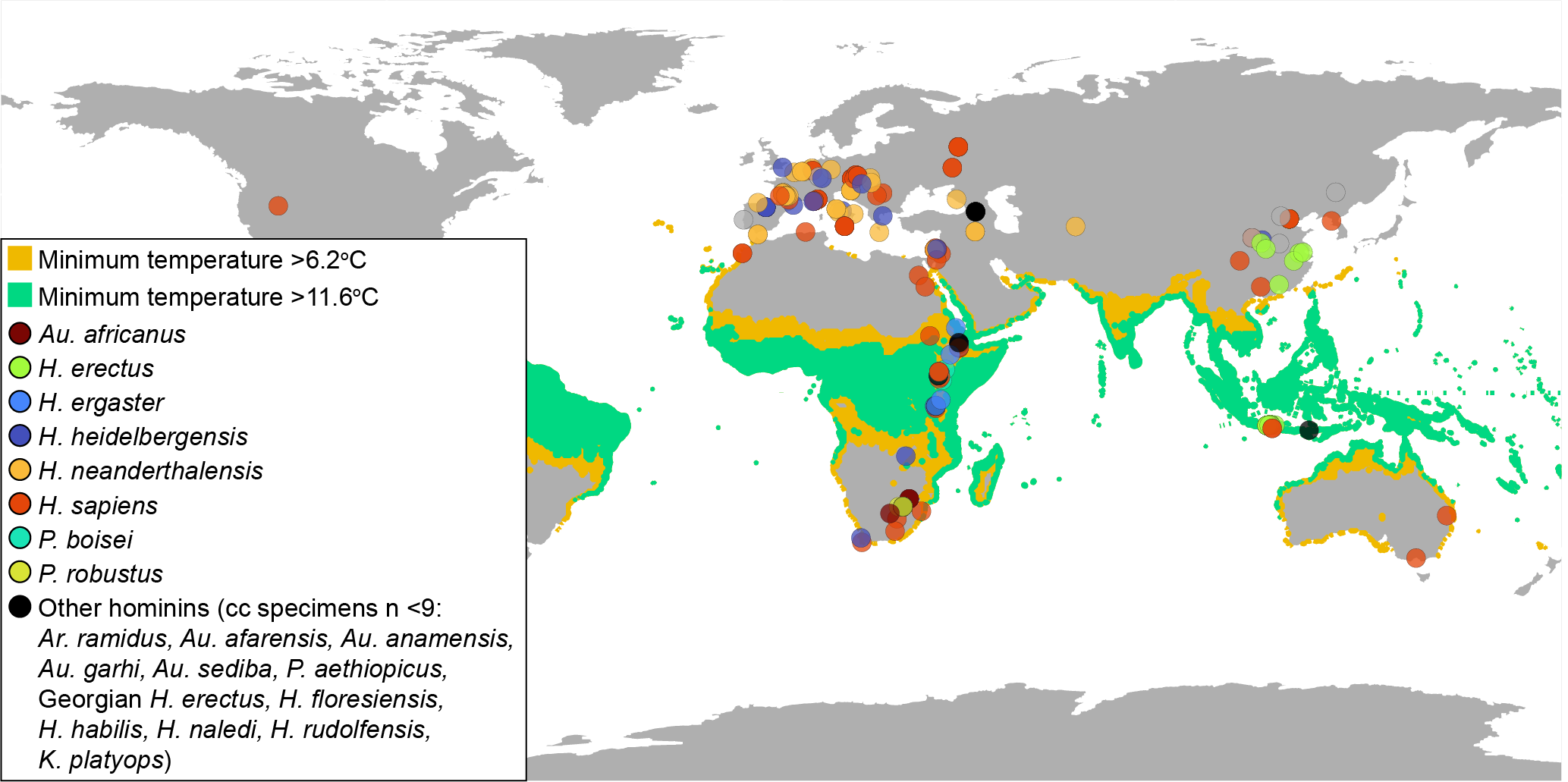

*Fig.S2. The lowest minimum daily temperatures recorded from 1950-2023 were plotted with fossil hominin cranial specimens. Daily temperature estimates were compiled from the KNMI climate explorer. This is not taken to represent the minimum possible temperature of those specimens during their lifetime, but to show 1) the spatial distribution of minimum temperatures that have occurred over a 73-year period (which cannot be accurately modelled by the PALEO-PGEM climate emulator) and 2) that much of Earth experiences temperatures that fall below the MSTs of H. erectus.*

### Glacial-interglacial variability

The Pleistocene is characterised by northern hemisphere glaciation, global cooling under a backdrop of increased lengthening and amplitude of glaciations. This contrasts the relatively warm and stable conditions of the preceding Pliocene. A transition from the so-called “40-kyr world” (where climatic variation was underpinned by low amplitude ~41-kyr cyclicity) to the “100-kyr world” (where amplitude increased yet frequency reduced to ~100-kyr cycles) occurred between 1.2-0.7 Ma. An additional increase in amplitude is considered to have taken place at ~460 ka, following the so-called “Mid-Bruhnes Event”. These changes in major global climate dynamics, particularly the dramatic fluctuations from globally wetter-warmer to drier-colder conditions, have routinely been proposed to have induced cultural, biological, and technological adaptation in hominins(10). The CoV of bio-1 (Mean Annual Temperature), bio-12 (annual rainfall) and NPP over 100 kyr eccentricity (CoV_100-kyr_) and 41 kyr obliquity (CoV_41-kyr_) cycles were run as independent test of the effect of long-term orbital-scale glacial-interglacial variability on encephalisation. All tests followed the aforementioned structure, including between- and within-species body mass, between- and within-species age and between- and within-species of the variable, whilst simultaneously accounting for phylogenetic effects and including a random effect of within-species body mass, within-species age and within-species of the variable in question. We found a significant effect of within-species Bio-1 CoV_100-kyr_, suggesting that interglacial-glacial variability of temperature has an influence on relative cranial capacity. As an additional check, we then tested this whilst including the within- and between-species effect of PC1. Together, the effects (between- and within-species) of PC1 and Bio-1 CoV_100-kyr_ are cancelled out and they both become insignificant. Indeed, these variables are highly correlated (0.75) suggesting that variance inflation leads to insignificant results. Such a high correlation to PC1 indicates that colder and more variable environments are subjected to greater variability on 100-kyr eccentricity cycles and thus these variables cannot be effectively distinguished.

| **Test variable** | **Description** | **Intercept** | **Between-species var slope** | **Between-species var pMCMC** | **Within-species var slope** | **Within-species pMCMC** | **R2 conditional** | **R2 marginal** | **h^2^** |
| --- | --- | --- | --- | --- | --- | --- | --- | --- | --- |
| CoV Bio-12 _41-kyr_ | 41 kyr moving coefficient of variation of annual precipitation | *-0.39* | *0.00* | *0.06* | *-0.03* | *0.12* | *0.95* | *0.73* | *0.85* |
| CoV Bio-12 _100-kyr_ | 100 kyr moving coefficient of variation of annual precipitation | *-0.41* | *0.00* | *0.09* | *-0.02* | *0.15* | *0.95* | *0.71* | *0.86* |
| CoV NPP _41-kyr_ | 41 kyr moving coefficient of variation of NPP | *-0.59* | *0.04* | *0.25* | *0.02* | *0.18* | *0.95* | *0.67* | *0.87* |
| CoV NPP _100-kyr_ | 100 kyr moving coefficient of variation of NPP | *-0.64* | *-0.01* | *0.37* | *0.02* | *0.14* | *0.95* | *0.65* | *0.88* |
| CoV Bio-1_41-kyr_ | 41 kyr moving coefficient of variation of mean annual temperature | *-0.67* | *-0.04* | *0.32* | *0.03* | *0.09* | *0.98* | *0.65* | *0.88* |
| **CoV Bio-1 _100-kyr_** | **100 kyr moving coefficient of variation of mean annual temperature** | ***-0.64*** | ***0.03*** | ***0.34*** | ***0.04*** | ***0.02*** | ***0.98*** | ***0.66*** | ***0.88*** |
| PC1 and CoV Bio-1 _100-kyr_† | Combination of PC1 and Bio-1 CoV_100-kyr_ | *-0.58* | PC1: *0.04*  Bio-1 CoV_100-kyr_:  *-0.09* | PC1: *0.31*  CoV_ecc_ Bio-1: *0.32* | PC1: *0.01*  Bio-1 CoV_100-kyr_: *0.03* | PC1: *0.21*  Bio-1 CoV_100-kyr_: *0.16* | *0.96* | *0.66* | *0.88* |
| Latitude | *Relative latitude (distance [^o^] from the equator)* | *-0.54* | *0.06* | *0.32* | *0.04* | *0.05** | *0.94* | *0.67* | *0.87* |
| G-MAT | *Global Mean Annual Temperature calculated from PALEO-PGEM* | *-0.61* | *-0.06* | *0.24* | *0.00* | *0.27* | *0.94* | *0.67* | *0.87* |
| G-MAT_LR04_ | *Global Mean Annual Temperature inferred from the LR04 δ^18^O_benthic_ global ice-volume stack* | *-0.69* | *0.17* | *0.24* | *0.00* | *0.27* | *0.94* | *0.67* | *0.87* |

*Tab. S3. Table of additional parameters tested individually. * denotes a p-value approaching, or close to, 0.05. † for these tests, the structure follows: Cranial capacity ~ species mean body mass: within species body mass: species mean age: within species age: species mean test1: within species test_1_: species mean test_2_: within species test_2_.*

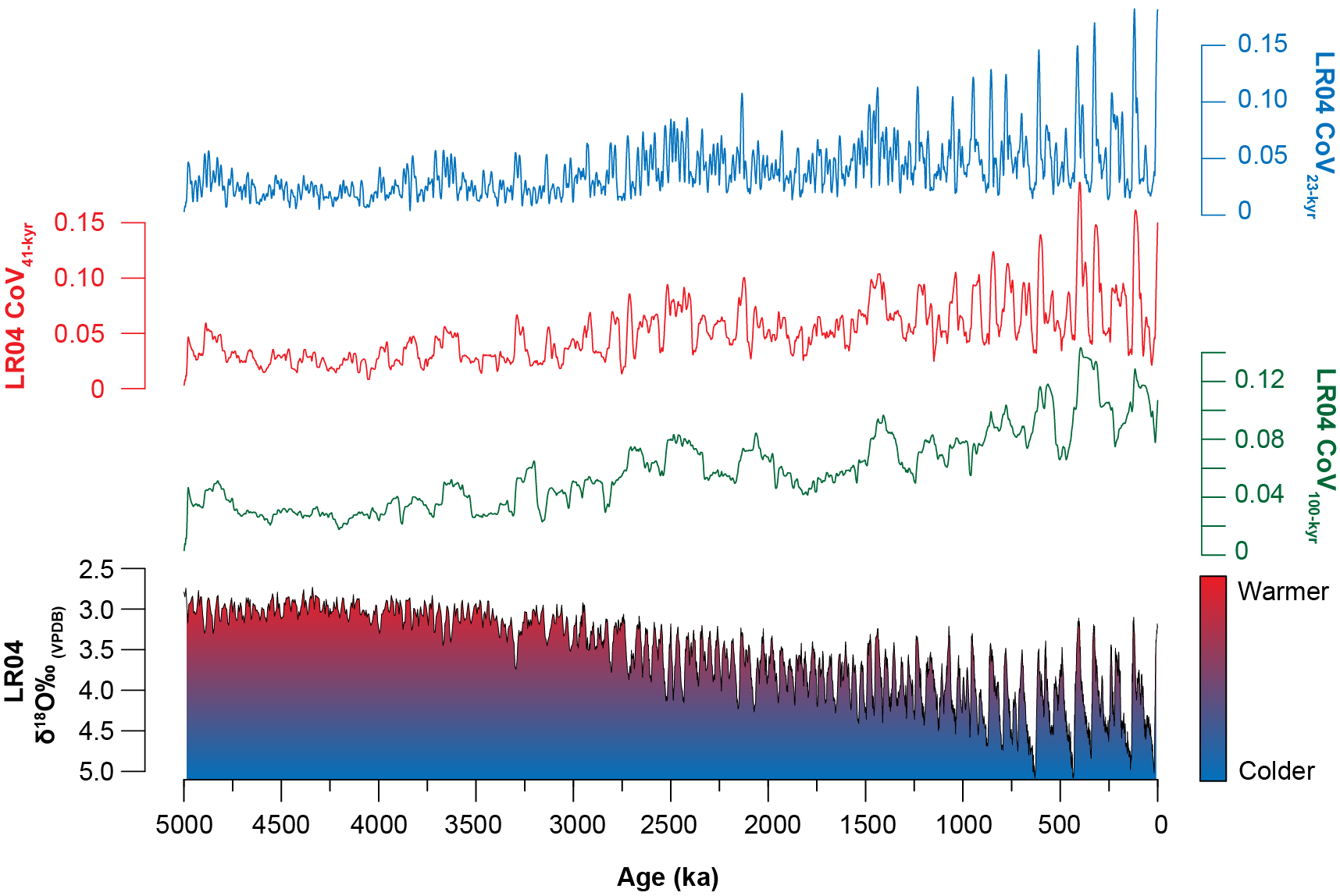

*Fig. S3. Representation of increasing interglacial-glacial variability over the last 5 Ma using the Coefficient of Variation (CoV) of the LR04 δ^18^O_benthic_ global ice-volume stack* (11) *on orbital 100-kyr (eccentricity), 41-kyr (obliquity) and 23-kyr (long precession) cycles.*
